# The cationic amino acid exporter Slc7a7 is vital for and induced in tissue macrophages with sustained efferocytic activity

**DOI:** 10.1101/2020.04.20.051664

**Authors:** Doris Lou Demy, Mireille Carrère, Ramil Noche, Muriel Tauzin, Marion Le Bris, Chooyoung Baek, Malika Yousfi, Ignaty Leshchiner, Wolfram Goessling, Philippe Herbomel

## Abstract

Most tissues harbor a substantial population of resident macrophages. It is not quite known yet how their quite diverse phenotypes are shaped by the functions that they assume in each tissue. In this study, we elucidate a functional link between the Slc7a7 cationic amino acid transporter and tissue macrophages. We had identified a mutant zebrafish devoid of microglia due to a mutation in the *slc7a7* gene. We found that in Slc7a7 deficient larvae, macrophages do enter the retina and brain to become microglia, but then die during the developmental wave of neuronal apoptosis, which triggers intense efferocytic work from them. A similar macrophage demise occurs at other tissues and stages whereby macrophages have to engulf many cell corpses, be it due to developmental or experimentally triggered cell death. We found that *slc7a7* is by far the main cationic amino acid transporter gene expressed in macrophages of wild type zebrafish larvae, and that its expression is induced in tissue macrophages within 1-2 hrs upon efferocytosis. Our data altogether indicate that a high level of Slc7a7 is vital not only for microglia but also for any steadily efferocytic tissue macrophages, and that *slc7a7* gene induction is one of the adaptive responses that allow them to cope with the catabolism of numerous dead cells without compromising their own viability.

## INTRODUCTION

Most tissues in vertebrates harbor resident macrophages. They are thought to be important for tissue homeostasis (Okabe and Medzhitov, 2016). They display quite diverse phenotypic traits among different tissues - which historically led to give them different names, e.g. “microglia” for the resident macrophages of the central nervous system (CNS). Many of these tissue resident macrophages do not derive from monocytes produced in the bone marrow. Instead, they originate from the yolk sac, and migrate to colonize embryonic tissues where they are ultimately maintained or self-renew throughout life (Perdiguero and Geissmann, 2016).

In the past years, several studies aimed at obtaining a more comprehensive view of the molecular requirements for the early establishment of microglia and tissue-resident macrophage populations in general, using the zebrafish model. Thus, the CSF-1 receptor and its ligand IL34 were found essential for the attraction of yolk sac derived macrophages to the cephalic tissues, and the subsequent establishment of primitive microglia as well as epidermal macrophages (Herbomel et al., 2001; Wu et al., 2018). In addition, developmental neuronal cell death was found to be important as well for microglia establishment in the midbrain optic tectum (Xu et al., 2016). In parallel, we and others performed forward genetic screens for zebrafish larvae devoid of microglia (Demy et al., 2017; Meireles et al., 2014; Rossi et al., 2015; Shiau et al., 2013) - based on the convenient staining of microglia in live zebrafish larvae with neutral red that we had introduced (Herbomel et al., 2001). The mutations causing microglia absence in the resulting mutants published so far all affected microglia or its precursors cell-autonomously, and encoded proteins with very diverse functions: a Nod-like cytoplasmic receptor (Nlrc3l/Nlrp12); a transmembrane phosphate exporter (Xpr1b); a nuclear transcription regulator (Trim33); and a transmembrane cationic amino acid transporter (Slc7a7). For most of them, it is still unclear how the causative mutation led to microglia absence. We originally found the *slc7a7* mutant in a screen performed together with F. Peri’s team, who have then published their study, which concluded that Slc7a7 identifies microglial precursors prior to entry into the brain (Rossi et al., 2015) and microglia thereafter. While our independent mapping of the mutation agreed on the nature of the causative mutation, our conclusions on the functional significance of Slc7a7 in microglia development are quite different. Therefore we present here our own study. In short, we show that *slc7a7* expression is induced in any efferocytic tissue macrophages in the zebrafish larva, and is vital for them whenever they have to provide a high rate of efferocytic activity. Our study also sheds light on how microglia acquires some of its long-term specific molecular signature.

## RESULTS

In a F3 screen following ENU-based mutagenesis, we identified *cerise*^*NO067*^ – hereafter referred to as *cerise* - as a recessive mutant that displayed no microglia vitally stained by Neutral Red (NR) at 4 dpf (Fig. 1A,B). *Cerise* mutants are still devoid of microglia by 9 dpf (Fig. 1C) but otherwise properly developed, with a swim bladder and robust blood circulation (Fig. S1A,B), and can survive up to 17 days (data not shown).

**Fig. 1.**
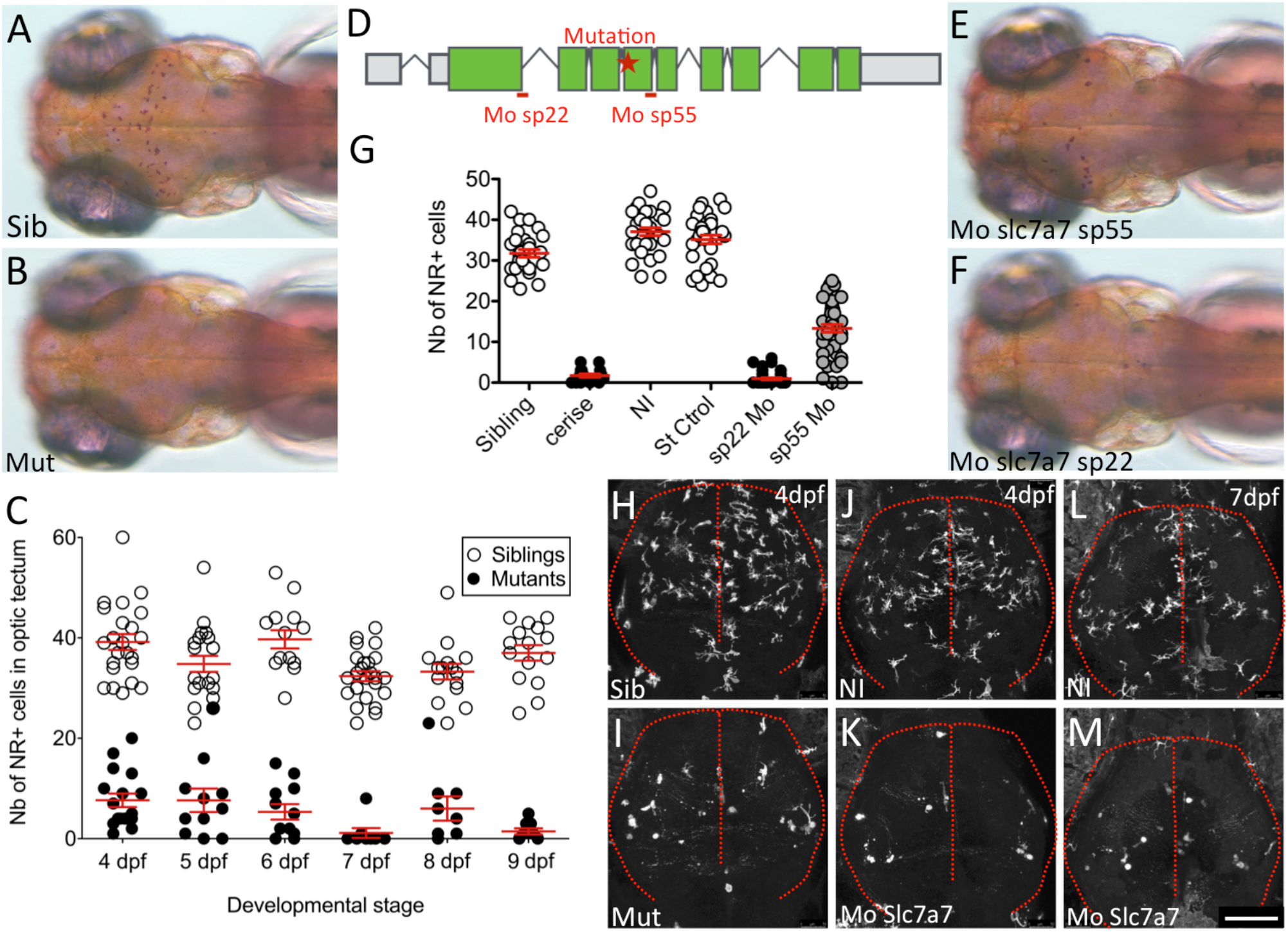
The *cerise* phenotype of microglia absence is caused by a mutation in the *slc7a7* gene and is fully mimicked by the sp22 morpholino. **(A, B)** Dorsal view of the microglia in the optic tectum of live 4 dpf larvae stained with Neutral Red (NR) in siblings **(A)** and cerise mutants **(B).** **(C)** Quantification of NR+ tectal microglia in *cerise* sibling (white dots) and mutant (black dots) larvae from 4 to 9 dpf. **(D)** Exon structure (green boxes) of the *slc7a7* gene, with position of the *cerise* mutation (red star) and of the splice junctions targeted by the sp22 and sp55 morpholinos (red dashes). **(E, F)** Dorsal view of the tectal microglia in live NR-stained sp55 (E) and sp22 (F) morphant larvae at 4 dpf. **(G)** NR+ tectal microglia cell counts show that unlike sp55, the sp22 morpholino fully mimicks the absence of microglia of *cerise* mutant larvae. **(H-M)** Dorsal view of mCherry+ tectal microglia in *Tg(mfap4:mCherryF)* zebrafish larvae at 4 dpf in control siblings (H), *cerise* mutants (I), uninjected controls (J), sp22 morphants (K), and at 7 dpf in uninjected controls (L) and sp22 morphants (M). Scale bar, 100 μm. Sib, sibling; mut, mutant; NI, non-injected; dpf, days post-fertilization; Mo, morpholino; NR+, Neutral Red positive; Nb, number

We first mapped the *cerise* mutation by bulk segregant analysis of SSLPs (Geisler et al., 2007) to a 7.7 Mb interval on linkage group 7 (LG7). We then moved to SNP analysis based on whole-genome sequencing (Leshchiner et al., 2012) of pools of mutant and sibling embryos (3 pools: 20 siblings, 20 mutants, then 200 mutants). This analysis refined the location of the causative mutation within the previously defined region of LG7, and led to identify in the mutants a homozygous A>T transversion at the splice junction between intron 4 and exon 5 of the gene *solute carrier family 7, member 7* (*slc7a7*) gene (Fig. 1D), confirming the results obtained by Rossi et al. (Rossi et al., 2015) in their independent characterization of the *cerise* mutant. This gene encodes the y+LAT1 chain of the heterodimeric transmembrane y+L cationic amino acid transporter (Torrents et al., 1999).

We evaluated the expression in macrophages/microglia of *slc7a7* and of the other genes encoding transmembrane cationic amino acid transport systems expressed at the plasma membrane, through an RNAseq analysis that we performed on fluorescent macrophages/microglia sorted from zebrafish larvae expressing mCherry under the control of a macrophage specific promoter (*mpeg1*) at 3 dpf (Fig. S2 and Table S1). We found *slc7a7* to be expressed at a high level in these cells, e.g. twice as much as the transmembrane receptor and macrophage lineage marker *csf1r*. *Slc3a2*, which encodes the CD98/4F2hc glycoprotein that heterodimerizes with Slc7a7 to constitute the y+LAT-1 transporter, was expressed at an even higher level (Table S1). In contrast, *slc7a6*, encoding y+LAT-2, the alternative possible partner of CD98/4F2hc in making a heterodimeric transporter, was expressed very weakly. Finally, of the three genes encoding monomeric cationic amino acid transporters - CAT-1 (*slc7a1*), CAT-2 (*slc7a2*) and CAT-3 (*slc7a3*) - only *slc7a2* and *slc7a3* were found expressed, but at a level 10-fold lower than *slc7a7* (Fig. S2 and Table S1). Thus, y+LAT1/Slc7a7 is the main cationic amino acid transporter in macrophages / microglia of zebrafish larvae.

To confirm that the *cerise* phenotype was caused by Slc7a7 deficiency and have an alternative tool to knock this gene down, we designed two different splice blocking morpholinos (Mo) (Fig. 1D). Upon injection into wild-type (wt) embryos, the effect of the sp55 morpholino - closest to the mutation, and used by Rossi et al. (Rossi et al., 2015) - on microglia was only partial, whereas the sp22 morpholino fully phenocopied the *cerise* mutation (Fig. 1E-G). Accordingly, subsequent experiments were conducted using the sp22 Mo.

Transferring the *cerise* mutation into the transgenic *Tg(mfap4:mCherryF)* background, which specifically highlights macrophages/microglia confirmed the lack of primitive microglia from 4 dpf onwards (Fig. 1H,I and Fig. S1C,D). The same was found in slc7a7-sp22 morphants (Fig. 1J-M). At earlier stages (2 dpf), Slc7a7 deficient embryos display a normal production and overall deployment of primitive macrophages and are phenotypically unrecognizable from their controls (Fig. S1E,F).

### Slc7a7-deficient macrophages die after colonizing the brain and retina

We noticed that the few microglial cells occasionally seen in the optic tectum (OT) of the mutants at 4 dpf, and even more so at 3 dpf (when the tectal microglia is just being established in the wt – see below), were more heterogeneous in size than wt microglia, and quite often occurred in small groups at two locations, near the midline and at the ventro-lateral corners of the OT (Fig. S3A-D) (this trait, not seen in other microglia mutants of our collection, is actually why we initially named this mutant *cerise*). Closer examination by VE-DIC microscopy revealed unusual unstained large vacuoles adjacent to or including the NR stained material (Fig. S3E-G), that have not been seen in wt larvae. Similar vacuoles appeared to be fluorescent in *cerise Tg(mfap4:mCherryF)* larvae, suggesting that they were remnants of dead/dying mCherry-positive macrophages/microglia (Fig. S3H-H’’). As macrophage death in the CNS or elsewhere cannot be assessed by the usual markers of apoptosis (e.g. annexin V staining, caspase3 activation, Tunel) because most of them contain engulfed dead cells that stain positive for all these markers, we turned to *in vivo* time-lapse confocal imaging. We followed the behaviour of fluorescent tectal macrophage/microglial cells in transgenic morphant larvae from 3 dpf onwards and compared it to that in control larvae (Fig. 2A-C, Movies 1 and 2). While the latter displayed the typical ramified and dynamic morphology with little movement of the cell center acquired by that developmental stage (Li et al., 2012; Peri and Nüsslein-Volhard, 2008; Svahn et al., 2013), their counterpart in the morphant still displayed a more macrophage-like morphodynamics (Movies 1 and 2). Then, over the next 10 hrs, they disappeared from the tectum (Fig. 2A-C) - mostly after rounding up, becoming inert, and being engulfed by another macrophage/microglial cell, which a moment later also underwent the same fatal fate (Fig. 2C and Movie 2). These observations indicated that Slc7a7-deficient macrophages were actually able to colonize the OT but died once there, which could explain their inability to ever establish as a proper microglial population.

**Fig. 2.**
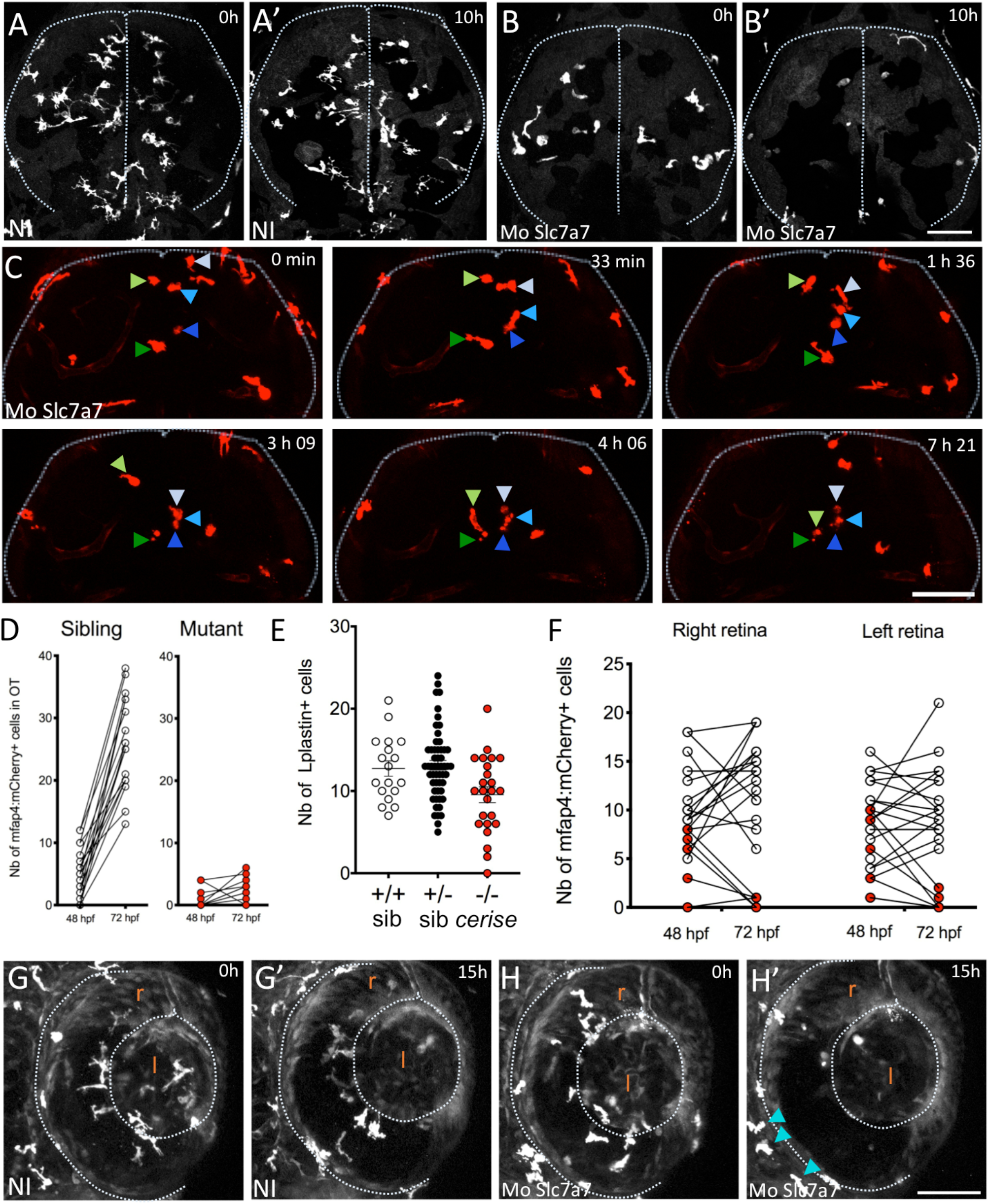
Macrophages/microglia in Slc7a7 deficient larvae die within the brain and retina from 2.5 dpf onwards. **(A-B’)** Dorsal view of the tectal microglia in live *Tg(mpeg1:Gal4; UAS:Kaede)* zebrafish larvae at the beginning (A, B) and end (A’, B’) of a 10 hours time-lapse confocal imaging session started at 72 hpf of a non-injected control (A, A’) and a sp22 morphant (B, B’). **(C)** Selected time points (maximum projections) of the *in vivo* time-lapse confocal imaging of Tg(*mpeg1:Gal4; UAS:NfsB-mCherry)* tectal microglia started at 72 hpf of a sp22 morphant, dorsal view. Grey or blue arrowheads, and green arrowheads, point at two series of macrophages that die, are engulfed by another, which dies in turn, and so on…; see also Movie 2. **(D)** *In vivo* quantification of mCherry+ macrophages/microglia in the retinas of *Tg(mfap4:mCherryF) cerise* siblings (white dots) and mutants (red dots) examined sequentially at 48 and 72 hpf. **(E)** Quantification of L-plastin+ macrophages in the retinas of *cerise* progeny at 42 hpf, subsequently genotyped as homozygous WT (+/+), heterozygous (+/−) or homozygous *cerise* mutants (−/−). **(F)** *In vivo* quantification of mCherry+ macrophages/microglia in the optic tectum of *Tg(mfap4:mCherryF) cerise* siblings (white dots) and mutants (red dots) examined sequentially at 48 and 72 hpf. **(G-H’)** Lateral view of the retinal microglia of live *Tg(mpeg1:Gal4; UAS:NfsB-mCherry)* larvae at the beginning (G,H) and end (G’, H’) of a 15 hours time-lapse imaging session started at 54 hpf of a non-injected control (G,G’) and a sp22 morphant (H,H’). See also Movies 3 and 4. Scale bars, 75 μm. r, retina; l, lens; NI, non-injected; h, hours; Mo, morpholino; Nb, number

To further explore this prospect of a transient colonization of the CNS in Slc7a7-deficient embryos, we aimed at examining earlier developmental stages. The OT and the retina are the two CNS structures most densely populated with microglia in zebrafish larvae, but they acquire most of it at different times. While the main wave of OT colonization occurs between 2.5 and 3.5 dpf ((Herbomel et al., 2001; Xu et al., 2016); see also Fig. 3D), the retinal microglia number has already reached its plateau by 48 hpf (Demy et al., 2017; Herbomel et al., 2001). We therefore counted the number of macrophage/microglia cells in the retinae by 42 hpf in the progeny of *cerise* heterozygote carrier fish, and then genotyped the embryos. The results demonstrated that macrophages in homozygous mutant embryos were indeed able to colonize the retinas quite efficiently (Fig. 2E). An independent live follow-up of individual embryos documented the subsequent decrease of the retinal macrophage/microglia population in the mutants between 2 and 3 dpf (Fig. 2F). Following our previous observations in the OT, we hypothesized that these retinal macrophages also died. So we time-lapse imaged the retinal macrophages in morphant vs. control embryos from 48 hpf onwards. In the time-lapse sequences shown in Fig. 2G-H’’ and Movies 3 and 4, while the retinal macrophage number in the morphant was similar to the control at the onset of the imaging, it then steadily decreased, to virtually zero by 15 hrs later, with evidence of macrophage rounding and demise (Fig. S4A-A’’). Interestingly, macrophages in interstitial tissues around the eye showed no sign of cell death (Fig. 2H’, cyan arrowheads).

**Fig. 3.**
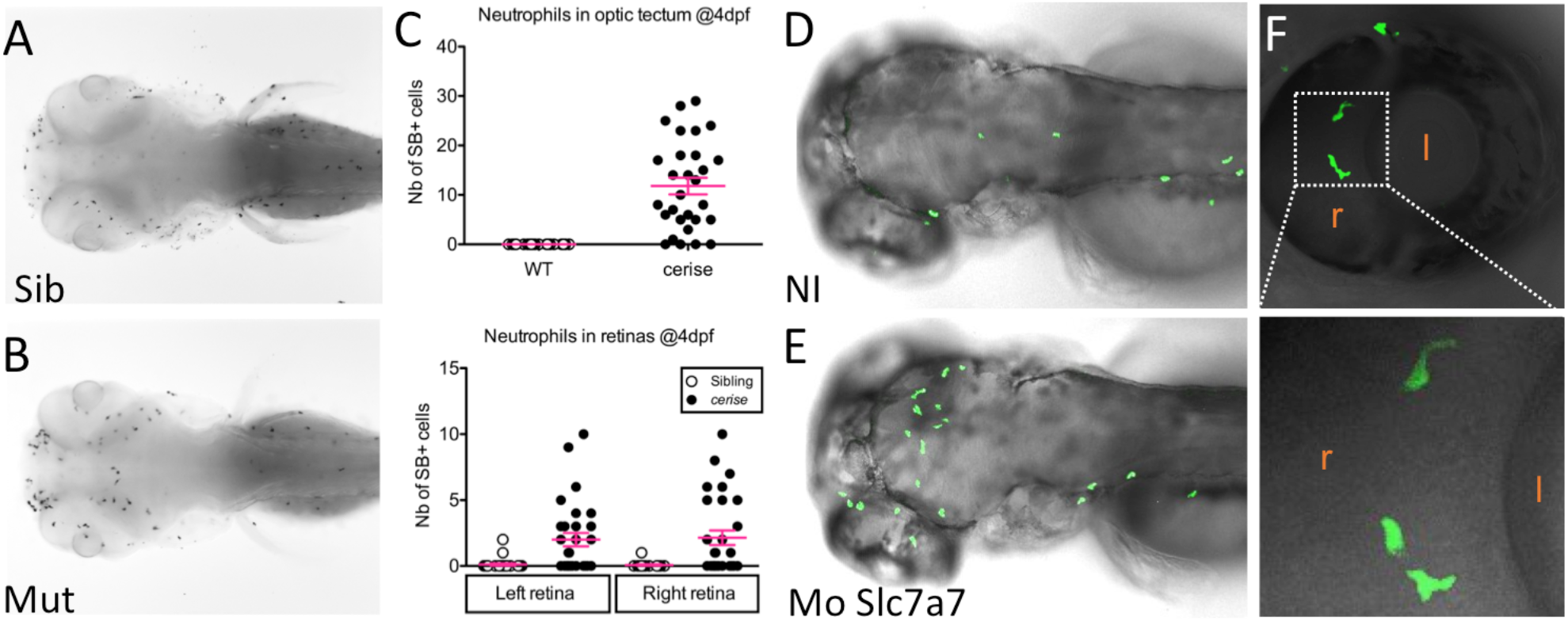
Neutrophils are recruited to the CNS of Slc7a7 deficient larvae following macrophage/microglia death. **(A-C)** Dorsal view of control **(A)** and *cerise* **(B)** larvae at 4 dpf stained for neutrophils with Sudan Black (SB), and corresponding SB+ cell counts in the optic tectum and retinas. **(C)** **(D-F)** Dorso-lateral view of live *Tg(mpx:GFP)* control **(D)** and *slc7a7* sp22 morphant **(E)** larvae at 3 dpf, and close-ups on GFP+ cells within the morphant retina at 3 dpf **(F**). See also Movie 5.

Thus, it appears that Slc7a7-deficient macrophages are able to normally colonize the OT and retina, but die there starting from 2.5 dpf. In the retina, where the microglial population has already reached its plateau by this stage, it results in a drop in cell number (Fig. 2E-F); in the OT, as the colonization wave and disappearance are concomitant, the microglial population never becomes substantial (Fig. 2D).

### Neutrophils infiltrate the brain and retina of Slc7a7-deficient larvae following microglia demise

In the zebrafish embryo, beside macrophages, primitive myelopoiesis in the yolk sac also produces neutrophils. Unlike in mammals, these neutrophils then disperse, live and wander in the tissues like the macrophages, except for the CNS that they normally never enter (Le Guyader et al., 2008). However, in cerise/slc7a7 mutants and morphants, we found a massive infiltration of neutrophils in the brain OT and the retinas by 4 dpf (Fig. 3A-C). Through time-lapse confocal imaging of GFP+ neutrophils in *Tg(mpx:GFP)* larvae, we found that this influx of neutrophils into the retinae and brain appeared to start by 2.5-3 dpf, when the macrophages/microglial cells were dying there (Fig. 3D-F and Movie 5). Time-lapse imaging of double transgenic lines highlighting macrophages in red and neutrophils in green confirmed the link between macrophage state and neutrophil recruitment, as the latter repeatedly interacted with the few macrophages remaining in the *cerise* brain (Fig. S5). However, neither qPCR for the cytokines that might be released by the stressed/dying macrophages (IL1-ß, TNF-α, CXCL8), nor immunohistochemistry for IL1-ß (Vojtech et al., 2012), nor slc7a7 MO injection into *Tg(IL1-ß:GFP)* reporter embryos (Nguyen-Chi et al., 2014) allowed us to identify the molecular signals responsible for the attraction of neutrophils into the CNS of mutant larvae, leaving them yet to be identified (not shown).

### Slc7a7 is required for the survival of highly efferocytic tissue macrophages

Thus, microglial cells in *cerise* larvae begin to die by 2.5 dpf. This is also precisely the stage at which WT microglial cells become intensely vitally stained by neutral red (Herbomel et al., 2001). As a weak base, neutral red accumulates in the phagolysosomal compartment. In the primitive microglia it accumulates over time in the phagosomes containing apoptotic bodies that have become acidified (Fig. S6A-A’’), The onset of intense NR staining of microglia by 56-60 hpf (Herbomel et al., 2001) thus actually reflects the beginning of the major wave of neuronal developmental cell death (DCD) and the engulfment of the numerous resulting corpses by the young microglia. As microglial cells in *cerise* embryos begin to die by that same stage, we hypothesized that Slc7a7 could be vital for macrophages / microglia that have to eliminate numerous apoptotic cells. We therefore examined other tissue macrophage populations that face such a situation at some point in zebrafish development. Beside microglia, we previously noted the similarly intense but more transient neutral red staining of the macrophages that eliminate the remnants of the hatching gland by 3 dpf (Fig. 4A) (Herbomel et al., 2001), and the macrophages of the Caudal Hematopoietic Tissue (CHT, (Murayama et al., 2006)), which engulf many of the circulating primitive erythrocytes particularly from 6 to 12 dpf (Fig. 4B,C), as the latter become progressively replaced by erythrocytes produced from the definitive hematopoiesis. Therefore we examined these highly efferocytic macrophage populations in *cerise* mutants and their siblings. In the latter, at the hatching gland location at 3 dpf, WT mCherryF+ macrophages, despite a heavy efferocytic load, still showed numerous long pseudopodia /ramifications (Fig. 4D). In sharp contrast, in *cerise* mutants, no ramified macrophages could be seen, and the widely scattered mCherryF+ spots, mostly of subcellular size, were strongly suggestive of macrophage degeneration (Fig. 4E). Similarly, in the CHT, *cerise* mutants display an accumulation of subcellular sized mCherry+ spots and much fewer healthy macrophages than in WT controls (Fig. 4F-I). Strongly neutral red stained macrophages can also occur more occasionally earlier in the blood flow, notably in the yolk sac where macrophages engulf the variable number of primitive proerythroblasts that undergo apoptosis around the onset of blood circulation (Herbomel et al., 1999; Herbomel et al., 2001) (Fig. S6B-C). In *cerise* morphants by this stage, beside NR++ efferocytic macrophages similar to WT, we occasionally observed by VE-DIC microscopy similarly intense neutral red stained material, marking phagocytosed proerythroblasts, within or adjacent to a larger unstained vesicle merely containing small particles in brownian motion (Fig. S5E; Movies 6 and 7) – a figure never seen in WT embryos, but similar to those previously mentioned in the optic tectum at 3 dpf (Fig. S3G), which most likely represents a macrophage that engulfed proerythroblasts and then died.

**Fig. 4.**
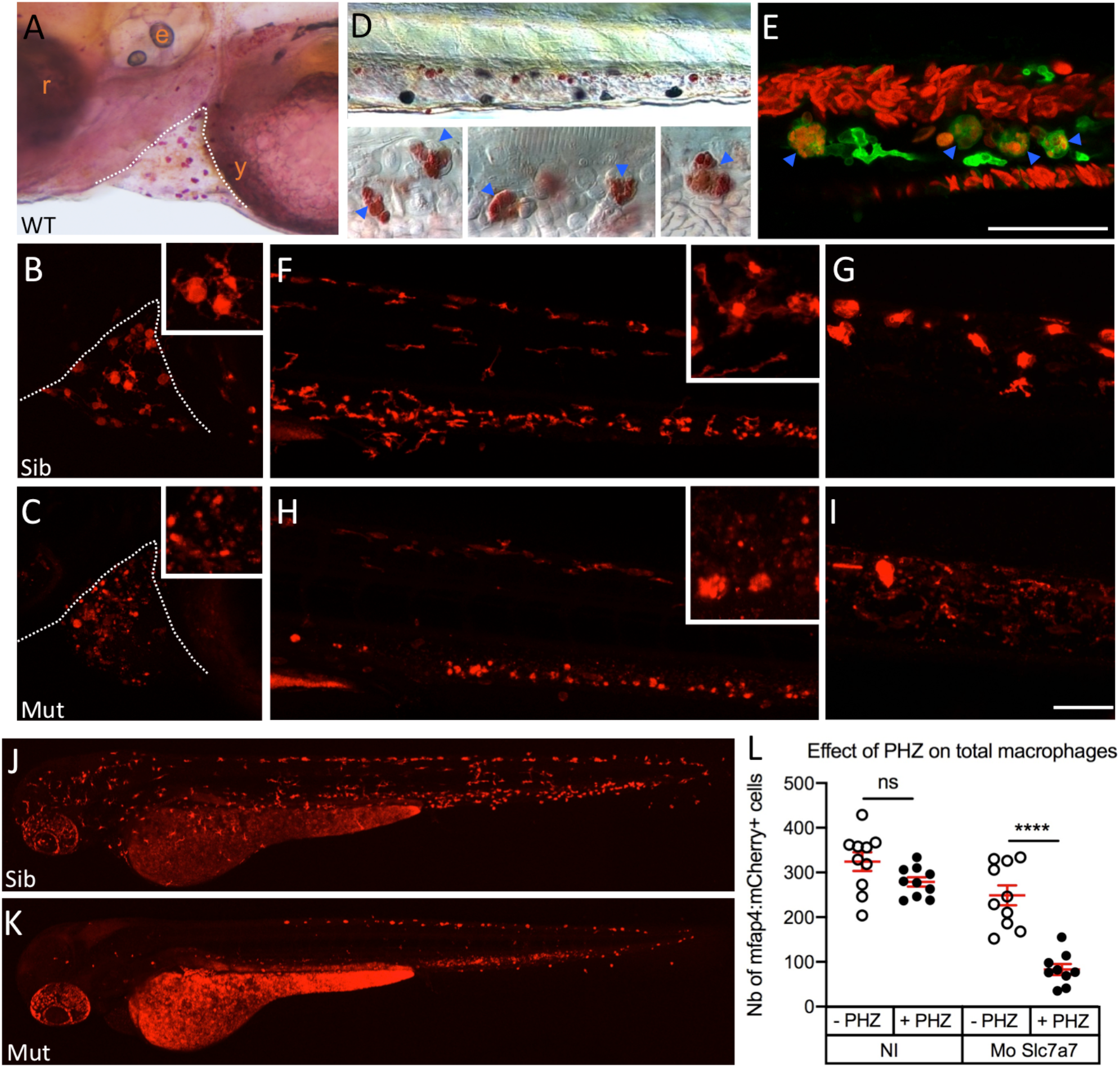
Tissue macrophages other than microglia also die upon sustained efferocytosis in *cerise* mutants. **(A-I)** Lateral view of the hatching gland **(A-C)** and CHT **(D-I)** in live WT larvae stained with Neutral Red at 3 dpf (A) and 7 dpf **(D)**, in *Tg[gata1:DsRed; mpeg1: GFP]* larvae at 7 dpf **(E),** and in sibling **(B, F, G)** and *cerise* **(C, H, I)** embryos carrying the *Tg[mfap4:mCherry]* or *Tg[mpeg1:mCherry]* transgene at 3 dpf **(B, C, F, H)** and 6 dpf **(G, I)**. **(J-L)** Lateral view of the global macrophage population at 54 hpf in live *Tg(mpeg1:mCherryF)* control **(J)** and *slc7a7* morphant **(K)** embryos treated with PHZ at 30 hpf for 20 hrs, and corresponding quantification **(L)**. Scale bars, 50 μm

These data collected across tissues and stages altogether strongly suggest that *slc7a7* expression is vital for any tissue resident macrophages that even transiently face the challenge of intense efferocytic work.

We further explored this hypothesis by inducing large-scale efferocytic work for macrophages in slc7a7 morphant vs. control embryos. To achieve this, we took advantage of the fact that blood circulation in zebrafish embryos is open until 2.5 dpf (due to its free flow in the yolk sac before the closure of the common cardinal vein) and thus readily accessible to all interstitial macrophages. We treated embryos at 30 hpf for 20 hrs with a low dose of phenylhydrazine (PHZ), which induces apoptosis of all circulating primitive erythrocytes (Ferri-Lagneau et al., 2012; Shafizadeh et al., 2004). Macrophages in slc7a7-sp22 morphants readily engulfed the apoptotic erythrocytes within a few hours as in WT control embryos, but on the next day, they had virtually disappeared from the slc7a7 morphant embryos treated with PHZ, while their number had only slightly decreased in the control embryos (Fig. 4J-L).

### Slc7a7 expression is induced in tissue macrophages upon efferocytosis

We next examined the expression of the *slc7a7* gene in wt larvae by whole-mount in situ hybridization (WISH) (Fig. 5). We noticed that the two tissue macrophage populations that showed a distinctly higher *slc7a7* expression by WISH were precisely the brain and retinal macrophages/microglia from 56 hpf onwards (i.e. the stage at which neuronal DCD begins) and the hatching gland associated macrophages by 3 dpf (Fig. 5A-C). From 3 dpf, again consistent with the death of Slc7a7-deficient macrophages, slc7a7 mRNA signal was also lightly detected in the tail (Fig. 5D), in addition to the kidney tubules and gut epithelium as in mammals (Fig. 5E).

**Fig. 5.**
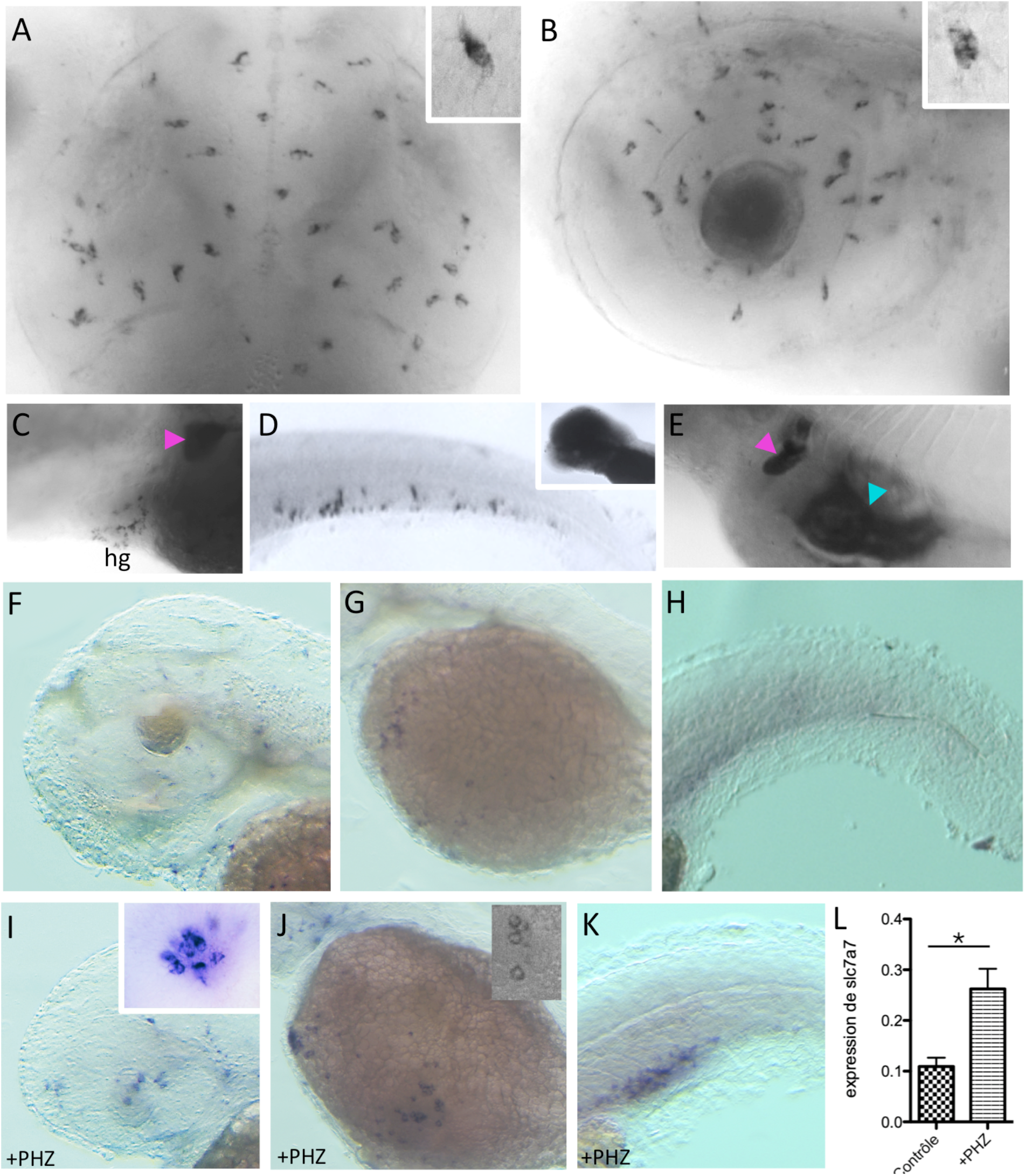
Efferocytic macrophage populations strongly express Slc7a7. Whole-mount *in situ* hybridization for *slc7a7* mRNA. **(A-E)** WT embryos **in** dorsal **(A)** and lateral **(B-E)** view. **(A)** Optic tectum, 3 dpf; **(B**) retina, 60 hpf; **(C)** hatching gland (hg), 3 dpf; **(D)** CHT, 3 dpf; this lower slc7a7 mRNA signal was seen when the anterior part was overexposed (inset). **(E)** pronepric tubules (pink arrowhead, also visible in C) and intestinal bulb (cyan arrowhead), 3 dpf. **(F-L)** Induction of slc7a7 expression in macrophages following PHZ treatment. Lateral views at 50 hpf in control (F-H) or PHZ treated (I-K) embryos. (F, I) Head, with close-up on slc7a7+ macrophages in hyaloid vasculature in I; (G, J) yolk sac, with close-up on slc7a7+ macrophages in the blood circulation valley (duct of Cuvier) in J; (H, K), tail, with slc7a7+ macrophages in the caudal vein plexus area in K. **(L)** Global quantification of slc7a7 expression by qPCR on control and PHZ-treated embryos. (*) p< 0.05.

We then examined *slc7a7* expression by WISH following PHZ-mediated death and engulfment of circulating erythrocytes, and found that it was again induced in the engulfing macrophages (Fig. 5F-K), enough to cause a 3-fold increase at the whole-embryo level (Fig. 5L). These results altogether indicate that *slc7a7* expression is not only required in macrophages with sustained efferocytic activity, but also induced by the latter, suggesting that a high level of Slc7a7 is necessary in this situation.

In order to refine the temporal link between efferocytosis and upregulation of Slc7a7 expression in macrophages, we aimed at inducing efferocytosis at precise anatomical locations and time. To do so, we exposed the fish embryos to a low concentration of CuSO_4_ (50 μM), which has been shown to rapidly and specifically induce apoptosis of the sensory cells of the neuromasts of the lateral line within the epidermis (Fig. 6A), causing the immediate recruitment of 5-6 macrophages per neuromast to engulf the dead cells (Carrillo et al., 2016; Olivari et al., 2008). We treated the fish embryos at 54 hpf for 30 min. and monitored *slc7a7* expression at each neuromast of the posterior lateral line by WISH every 30 min over the next 6 hours (Fig. 6B). The CuSO_4_ treatment efficiently destroyed the sensory cells (Fig. 6C-E), hence recruiting macrophages (Fig. 6C’-E’), which started to express *slc7a7* strongly enough to be detected by WISH (Fig. 6C’’-E’’,F,G), with a peak by 1.5 hr post treatment (hpt), and virtually no more expression by 4 hpt (Fig. 6B), even though recruited macrophages were still present in the area (see also Carrillo et al., 2016).

**Fig. 6.**
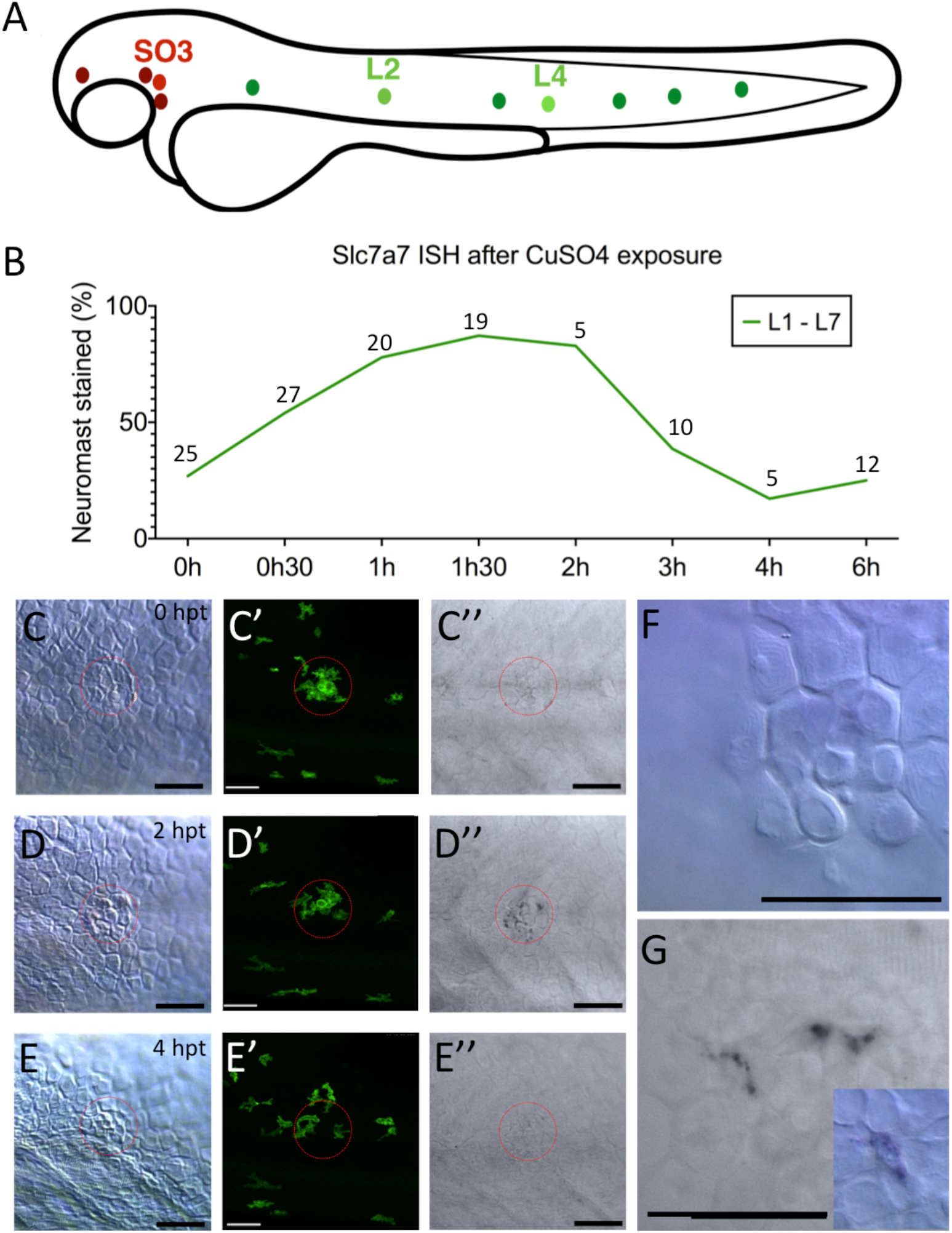
Slc7a7 expression is maximally induced 1-2 hrs following efferocytosis. (A) Location of the neuromasts of the posterior lateral line - named L1 to L7 in rostro-caudal direction - in the zebrafish embryo at 48 hpf. (B) Fraction of neuromasts showing a clear slc7a7 mRNA signal by whole-mount *in situ* hybridization at successive time points after a 30 min exposure to CuSO_4_. The number of embryos screened for each time point (n) is indicated above the curve. (C-G) VE-DIC/Nomarski (C,D,E,F), confocal (C’,D’,E’) and bright-field observation (C’’,D’’,E’’,G) of neuromast morphology, *Tg(mpeg1:GFP)* macrophage recruitment, and slc7a7 ISH signal at 0 (C-C’’), 2 (D-D’’) and 4 (E-E’’) hours after exposure to CuSO4. Close-up observation at neuromast L4 at 2 hpt shows that *slc7a7* is expressed by macrophages recruited at the damaged neuromast (F-G). Scale bars, 50 μm

Taken together, our results thus show that the cationic amino acid exporter Slc7a7 is vital for and induced in tissue macrophages with sustained efferocytic activity, rather than being a mere marker of microglia and its precursors.

### The 4C4 “microglia marker” is induced in highly efferocytic tissue macrophages

Another “microglia marker” has been the 4C4 monoclonal antibody, the only one so far available to specifically immunodetect microglia in the CNS of adult and larval zebrafish (Becker and Becker, 2001; Chia et al., 2019; Mazzolini et al., 2020; Ohnmacht et al., 2013; Tsarouchas et al., 2018). Yet when we examined the development of 4C4 immunoreactivity during zebrafish ontogeny, we found that it followed a pattern very similar to *slc7a7* expression. Firstly, it begins to label macrophages/microglia in the brain and retina by 2.5 dpf, i.e. the onset of the neuronal DCD wave and correlative heavy vital staining of CNS macrophages by Neutral Red (Fig. 7A,B); moreover, high magnification views showed that it especially decorated the most visibly efferocytic macrophages/microglial cells there (Fig. 7C). Secondly, by 3 dpf it labelled not only the microglia, but also, just as strongly, the efferocytic (NR++) macrophages eliminating the remnants of the hatching gland (Fig. 7D-E). Thirdly, in a serendipitous occurrence of embryo clutches in which 1/4 of the embryos displayed an abnormally early wave of apoptosis in the CNS by 35-48 hpf (Fig. 7F,G), we witnessed in the latter embryos a correlative influx of highly efferocytic (NR++) macrophages in the affected CNS areas (Fig. 7 H,I), and these macrophages were strongly 4C4-immunoreactive (Fig. 7 J,K).

**Fig. 7.**
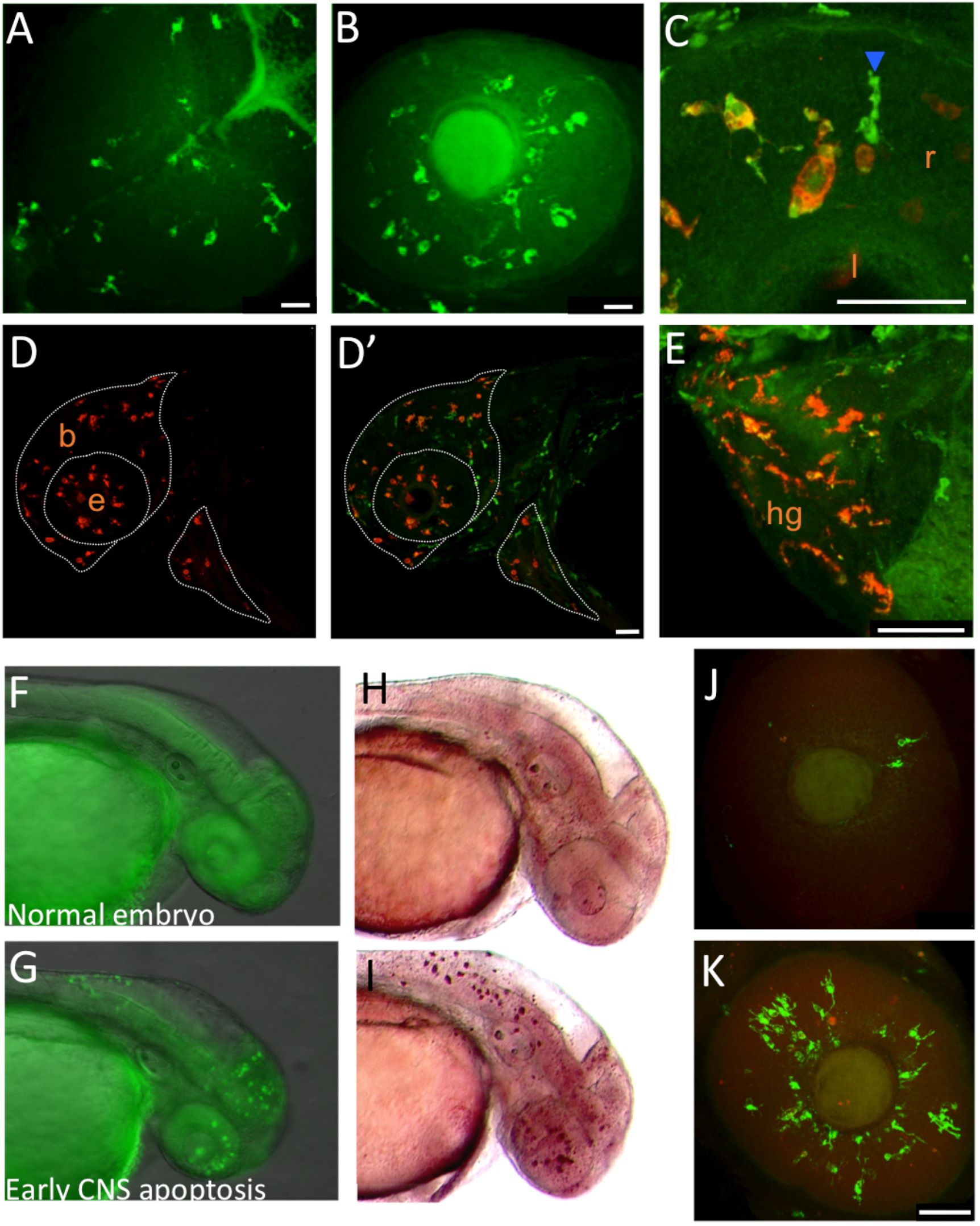
Expression of the 4C4 “microglia marker” is actually induced in tissue macrophages upon efferocytosis. (A, B) Dorsal view of the optic tectum (A) and lateral view of the retina (B) of 4C4 immunostained WT embryos at 60 hpf. (C-E) Lateral view of WT larvae at 72 hpf immunostained for 4C4 (red), and Pu.1 (green) which highlights leukocytes. (C) Close-up on the retina; one macrophage/microglia displaying no clear efferophagosome is Pu.1+, 4C4- (blue arrowhead) while two other microglial cells that display clear effero-phagosomes are Pu.1+, 4C4+ (white arrowheads). (D, D’) head and dorso-anterior yolk sac /hatching gland area. (E) close-up on the hatching gland area. (F-K) In a series of embryo batches that serendipitously contained 1/4 of embryos (G, I, K) displaying an abnormal wave of cell death by 35-48 hpf in the CNS, as shown by live acridine orange (AO) staining (F, G), numerous NR++ macrophages are present in the same CNS areas (H, I), and these macrophages are 4C4++ (J, K). Scale bars, 50 μm

Altogether these data clearly evidence that just like *slc7a7* expression, the unknown epitope recognized by the 4C4 antibody is actually induced in macrophages upon efferocytosis, and only later becomes stably expressed in the microglia up into adulthood.

## DISCUSSION

In this study, we have found that Slc7a7-deficient zebrafish lack microglia because Slc7a7 is vital for tissue macrophages with sustained efferocytic activity. Thus, macrophages that have entered the brain and retina to become microglia start to die when the main wave of neuronal DCD occurs, triggering intense efferocytic work from these macrophages to eliminate the numerous neuron corpses. In the optic tectum - the most routinely documented brain compartment in zebrafish microglia studies - the initial entry of Slc7A7-deficient macrophages can be easily overlooked because the main developmental influx of macrophages into the OT is triggered by signals emanating from the dying neurons (Xu et al., 2016). Therefore when the neuronal DCD wave begins by 2.5 dpf, the attracted macrophages immediately have to engulf many neuron corpses, and die right away if they are Slc7a7-deficient. In support of this scenario, we had initially noticed that the few neutral red labelled microglial spots found in the mutant OT, usually together with signs of microglial demise, often occured in the ventrolateral corners of the OT (hence the name *cerise).* A recent study precisely showed that these are indeed the main sites of macrophage entry into the OT (Xu et al., 2016).

In contrast, in the retina, the macrophage/microglia number has already reached its plateau by 2 dpf; therefore in Slc7a7-deficient embryos, the demise of this already established population when the neuronal DCD begins is much more evident than in the OT.

In both the OT and the retina, we found a substantial infiltration of neutrophils following the demise of macrophages / microglia there. This neutrophil infiltration cannot be merely due to the accumulation of neuronal corpses in absence of microglia to engulf them, since it does not occur in other microglia-deficient mutants such as the CSF1R-deficient *panther* (Herbomel et al., 2001; Wu et al., 2018). A similar neutrophil infiltration was reported in one particular mutant, deficient in Nlrc3l (Shiau et al., 2013), in which macrophages showed an early inflammatory phenotype, with a clear induction of *il1-ß* and *cxcl8*. In contrast, we found no induction of these cytokine genes in *cerise* embryos in toto, nor any sign of *il1-ß* induction in Slc7a7-deficient macrophages in situ using a Tg(il1-ß:GFP) reporter line (Nguyen-Chi et al., 2014). So the molecular signals attracting neutrophils to the dying microglia in the CNS remain to be identified.

Since our initial characterization of zebrafish primitive microglia (Herbomel et al., 2001), its intense vital staining by Neutral Red has been widely used as an easy readout for microglia number, notably in forward and reverse genetic screens (Kuil et al., 2019, and references therein). Here we clarify that this strong NR staining actually reflects the specific accumulation of neutral red over time in the already acidified phagosomes containing apoptotic bodies. Therefore beyond microglia, it is a very convenient live staining method to spot any episode of high efferocytic activity among the various tissue macrophage populations of the developing fish. We have thus identified two such episodes, and the correlative demise of the corresponding macrophage populations in the *cerise* mutant – i) the engulfment by 3 dpf of the hatching gland cells that died following the release by apocrine secretion of their protease-loaded granules into the medium to trigger hatching; and ii) the engulfment of many primitive erythrocytes once the definitive ones are produced, peaking from 5-6 dpf onwards. These examples further indicate that Slc7a7 is vital not specifically for microglia but for all tissue macrophages that undergo a phase of high efferocytic activity.

In line with these findings, we also found that Slc7a7 expression is induced in efferocytic macrophages in vivo, and our assay of macrophage recruitment to chemically damaged neuromasts showed that *slc7a7* induction occurs within 1-2 hrs following macrophage contact with apoptotic cells, and is over 2 hrs later. For macrophage populations whose efferocytic activity is sustained over many hours, or days as the primitive microglia, Slc7a7 expression becomes continually stimulated accordingly. It is conceivable that such continual stimulation may eventually turn into a stable expression feature of such a tissue macrophage population, even once their sustained efferocytic activity has subsided. We note indeed that in comparative transcriptome studies, *slc7a7* was found expressed at a high level in adult zebrafish microglia relative to brain tissue (Oosterhof et al., 2017), and in adult mouse microglia relative to peritoneal macrophages (Hickman et al., 2013). Similarly, we found here that the monoclonal antibody 4C4, which marks both resting and activated microglia throughout the CNS of adult and larval fish (Becker and Becker, 2001; Ohnmacht et al., 2013), actually labels tissue macrophages during zebrafish development wherever and whenever they are highly efferocytic – not only in the CNS. We found a similar developmental expression pattern for at least two other genes, coding for lysosomal proteins involved in protein and lipid catabolism, cathepsin B and prosaposin (P.H., unpublished data), which then will also remain highly expressed in adult microglia (Oosterhof et al., 2017). These observations open the path to a better understanding of how macrophages “differentiate” into microglia, and the biological mechanisms underlying the acquisition of their specific molecular signature. This is not to say that all microglia-specific traits will be acquired according to the above pattern. Notably, the expression of *apoeb* (encoding apolipoprotein E), also a marker of microglia in developing zebrafish (but not in adult mammals, in which it is expressed by many tissue macrophages), is not induced in efferocytic macrophages outside the CNS (Herbomel et al., 2001).

Thus our conclusion is quite different from that of Rossi et al. (Rossi et al., 2015) who claimed that Slc7a7 expression identifies microglial precursors prior to their entry into the brain. This conclusion was drawn from an embryo-wide photoconversion at 24 hpf of Kaede protein expressed from a transgenic slc7a7 locus; at 4 dpf they found photoconverted Kaede in microglial cells. However, the photoconverted Kaede in the microglial cells at 4 dpf appeared to delineate not the entire cells but an intracellular compartment - most likely phagolysosomal (Rossi et al., 2015, Fig. 4P-R), and the ~ 4 slc7a7+ macrophages that can be detected by 24 hpf (Rossi et al., 2015, Fig. 3K) could hardly account for the amount of photoconverted Kaede seen in most or all microglial cells at 4 dpf (Rossi et al., 2015, Fig. 4N). Our own interpretation of these data is that, owing to the high diffuse *slc7a7* expression seen throughout the head at 24 hpf (Rossi et al., 2015, Fig. 3I), the global photoconversion performed at 24 hpf notably photoconverted Kaede throughout the brain (Rossi et al., 2015, Fig. 4K,N), and this photoconverted Kaede accumulated in the phagolysosomal compartment of the microglial cells when the latter massively engulfed dead neurons during the neuronal DCD wave.

Why is Slc7a7 vital for highly efferocytic macrophages ? Sustained efferocytosis requires macrophages to catabolize the content of numerous cell corpses at a high rate, and then recycle or expell the resulting metabolites. These challenges involve adaptations of the macrophage metabolism that are only beginning to be uncovered (Han and Ravichandran, 2011). Since Slc7a7 is the main cationic amino acid transporter expressed in the macrophages, and is known to act best as an exporter, Slc7a7 deficiency would predictably lead to a large excess of cationic amino acids in efferocytic macrophages. An excess of arginine could conceivably be detrimental to the cell if it is used to produce excess NO via NO synthases (NOS); however we found that zebrafish primitive macrophages do not express any *nos* gene (Table S1). So the basis of a potential toxicity of excess intracellular cationic amino acids for the macrophage would remain to be identified.

Finally, could our results help understand some of the pathological outcomes of LPI (caused by inborn Slc7a7 deficiency (Torrents et al., 1999)) in humans ? Beside the obvious adverse consequences of Slc7a7 absence in the gut and in kidney tubules, LPI patients suffer from variable, quite diverse types of symptoms, at least two of which suggest an involvement of macrophages - pulmonary alveolar proteinosis, and hemophagocytic lymphohistiocytosis (Mauhin et al., 2017; Ogier de Baulny et al., 2012). It has been very difficult to study the etiology of these symptoms due the the lack of a viable mouse model until very recently (Bodoy et al., 2019). In addition to the symptoms listed above, a recent update of LPI patients follow-up pointed that over half of the patients presented with cognitive disorders (Mauhin et al., 2017). This has been explained so far by chronic or episodic hyperammonemia (due to a urea cycle disorder consequent to low serum arginine). Our study suggests an alternative or additional possible cause. As nothing is known about microglia in LPI patients, it is conceivable that as in the zebrafish, the first wave of microglia died in utero during the developmental neuronal death episodes, and that its demise contributed to later cognitive disorders.

## MATERIALS AND METHODS

### Zebrafish lines and embryos

Wild-type, transgenic and mutant zebrafish embryos were raised at 28°C in Embryo Water (Volvic© water containing 0.28 mg/mL Methylene Blue [M-4159; Sigma] and 0.03 mg/mL 1- phenyl-2-thiourea [P-7629; Sigma]), and staged according to Westerfield (Westerfield, 1993). AB & Tübingen wild-type (WT) fish (ZIRC), and the transgenic lines *Tg(mfap4:mCherry-F)^ump6^* (Phan et al., 2018), *Tg(mpeg1:Gal4FF)^gl25^* (Palha et al., 2013), *Tg(UAS:Kaede)rk8* (Hatta et al., 2006), *Tg(mpeg1:mCherry-F)^ump2^* (Nguyen-Chi et al., 2014), *Tg(UAS-E1b:Eco.NfsB-mCherry)^c264^* (Davison et al., 2007), *Tg(elavl3:EGFP)^knu3^* (Park et al., 2000), *Tg(mpx:GFP)^i114^* (Renshaw et al., 2006), *Tg(gata1:dsRed)^sd2^* (Traver et al., 2003), *Tg(mpeg:GFP-CAAX)^sh425^* (Keatinge et al., 2015), *Tg(lyz:DsRed2)^nz50^* (Hall et al., 2007) have been used in this study. The *cerise*^*NO067*^ mutant was obtained by ENU chemical mutagenesis on a Tübingen WT background, and maintained by outcross of heterozygous carriers with WT/Tg fish.

### Mapping and identification of the mutation

1500 *cerise* homozygous mutant larvae were collected along with a few of their siblings after neutral red based screening at 4 dpf. We first mapped the mutation to a 7.7 Mb interval on linkage group 7 (LG7) by bulk segregant analysis of sequence-length polymorphism (SSLP) markers, as described previously (Geisler et al., 2007). Then whole genome Illumina HiSeq2000 sequencing (Leshchiner et al., 2012) of 20 siblings and 220 mutants followed by SNP analysis led to the identification of a mutation within this region, a A>T change at the splice acceptor of *slc7a7* intron 4 (GenBank: BC110115.1).

### Morpholino injections

Two splice blocking antisense morpholinos (Mos) against slc7a7 RNA were synthesized by Gene Tools:

Sp22 (targeting the exon2/intron2 donor site): 5’-ATACATCCAACTCACAGATGCAGGC-3’

Sp55 (targeting the exon5/intron5 donor site): 5’-AAAGTGTTTATTACTCACCACAGCC-3’

1-5 nL of 0.6 mM Mo solution was microinjected in one to two-cell-stage embryos.

### Neutral Red vital staining of microglia and Sudan Black staining of neutrophils

Highly efferocytic cells, including microglia, were revealed in live embryos and larvae by adding Neutral Red [N-4638; Sigma] to Volvic Water at a final concentration of 5 μg/mL for 2 hrs, or 2.5 μg/ml for ~5 hrs when optimal signal-to-noise ratio was desired. After incubation in the dark at 28°C, larvae were rinsed in Volvic Water, anesthetized, and observed under a stereomicroscope or a compound wide-field microscope (see below). Sudan Black staining of neutrophils in formaldehyde fixed embryos was done as previously described (Le Guyader et al., 2008).

### Neuromast damage with CuSO4

Neuromast hair cell damage was achieved by incubating 54 hpf larvae in 50 μM CuSO4 [1.02791; Sigma] for 30 min, as previously described (Olivari et al., 2008). Treated larvae were then rinsed 3 times with Embryo Water. After WISH, *slc7a7* RNA signal was manually assessed for each neuromast (SO1-L7) using a Macrofluo microscope (Leica), and/or imaged on a Reichert Polyvar2 wide-field microscope (see below).

### Ablation of erythrocytes with Phenylhydrazine

The day of the experiment, Phenylhydrazine (PHZ) [P26252; Sigma] was dissolved in water at a concentration of 5 mg/mL for stock solution. Dechorionated live embryos at 30-35 hpf were then soaked in 5 μg/mL PHZ in Embryo Water overnight in the dark so as to induce massive death of primitive erythrocytes (Pelster and Burggren, 1996). The embryos were washed the next day with Volvic© water and processed for further experiments.

### Whole-mount mRNA in situ hybridization and immunohistochemistry

Embryos and larvae were anesthetized with 0.16 mg/ml Tricaine [A-5040; Sigma] at the stage of interest, then fixed overnight at 4°C in 4% methanol-free Formaldehyde [Polysciences, Cat#: 040181]. Whole-mount *in situ* hybridization (WISH) was performed according to (Thisse and Thisse, 2004). Whole-mount immunohistochemistry (WIHC) was performed as described previously (Murayama et al., 2006), omitting the acetone treatment. The primary antibodies used were rabbit anti-zebrafish L-plastin and PU.1 polyclonal antibodies (at 1:5000 and 1:800 dilution, respectively) (Le Guyader et al., 2008), and the 4C4 mouse monoclonal antibody (Becker & Becker, 2001), used as pure hybridoma supernatant. The secondary antibodies were Cy3-coupled anti-rabbit antibody [111-166-003; Jackson Immunoresearch] at 1:800 dilution, Cy3-coupled anti-mouse antibody [115-166-003; Jackson Immunoresearch] at 1:500 dilution, and Alexa488-coupled anti-rabbit antibody [A11070; Molecular Probes] at 1:500 dilution.

### SNP genotyping of fixed embryos

After WISH or WIHC and cell counting of macrophages / microglia cells in a clutch of *cerise* and sibling embryos, the tail tip of each embryo was cut using a sharp scalpel and isolated in a PCR tube for genotyping. Genomic DNA was extracted by boiling at 90°C for 30 min. in 40 μL of Extraction Buffer (25 mM NaOH, 0.2 mM EDTA), then the pH was neutralized by adding Tris-HCl. Genotyping was performed by qPCR using forward primers ending at the *cerise* mutation:

WT-complementary forward primer 5′-CCCTCTGGTGGTTATTTTTTCA-3′

mutant-complementary forward primer 5′-CCCTCTGGTGGTTATTTTTTCT-3′

The reverse primer 5′-CCACAGCATCACTGTCCAG-3′ was designed so as to generate a 142 nt amplicon, optimal for qPCR reaction. qRT-PCR was performed using Takyon Rox SYBR Master mix blue dTTP kit [Eurogentec, Cat#: UF-RSMT-B0701], and carried out for three biological replicates with measurements taken from three technical replicates on an Applied Biosystems 7300 Real Time PCR system (Thermofisher). Mutants and siblings were then sorted by comparing their amplification curves upon qPCR with the WT and mutant set of primers (Ct_mutant_ – Ct_WT_).

### Microscopy and image analysis

Low-magnification bright-field images were acquired using video cameras mounted on a Leica Macrofluo driven by the Metavue (Metamorph) software or a Zeiss Macrofluo driven by the Zen software (Zeiss).

Wide-field video-enhanced (VE) Nomarski / differential interference contrast (DIC) and fluorescence microscopy were performed as described previously (Herbomel & Levraud, 2004; Murayama et al., 2006), through the 40x/1.00 water-immersion objective of a Nikon 90i microscope and the 40x/1.00 oil-immersion objective of a Reichert Polyvar 2 microscope. Images were obtained from a HV-D20 3-CCD camera (Hitachi), digitized through a GVD-1000 DV tape recorder (Sony), then still images were collected using the BTVpro software (Bensoftware, London).

For fluorescence confocal imaging, embryos and larvae were mounted as previously described (Demy et al., 2017). Images were then captured at selected times on an inverted Leica SP8 set-up allowing multiple point acquisition, so as to image mutants and their siblings in parallel. Image stacks were processed with the LAS software to generate maximum intensity projections, or were exported into the Imaris software (Bitplane) for further analyses.

### RNAseq analysis of macrophages in zebrafish larvae

Zebrafish larvae from three different transgenic backgrounds highlighting macrophages - *Tg(mpeg1:mCherry)*, *Tg(mpeg1:Gal4; UAS:NfsB-mCherry)*, and *Tg(mpeg1:mCherry; mpx:gfp)* – were anesthesized at 3 dpf with Tricaine, then dissociated into single-cell suspensions as previously described (Covassin et al., 2006). The red fluorescent macrophages were sorted by FACS, collected directly in lysis buffer, and RNA was extracted using a miRNeasy Micro kit [217084; Qiagen]. Sequencing was performed on Illumina HiSeq2500 by ZF-Screens (Leiden, Netherlands) on three biological replicates: Tg(mpeg1:mCherry) (RNA extracted from 9480 cells of 92 embryos), Tg(mpeg1:Gal4/UAS:NfsB-mCherry) (RNA extracted from 10875 cells of 146 embryos), Tg(mpeg1:mCherry/mpx:gfp) (RNA extracted from 10004 cells of 196 embryos). 10 Mreads of 50 nt (0.5 Gb) were obtained for each replicate. The resulting data were analysed with the DESeq software.

## Acknowledgements

We wish to thank N. Trede and all members of the Trede lab for hosting and guiding D L Demy for the initial mapping of the *cerise* mutation, J-P Levraud for the mutant genotyping method, and our fish facility team for their excellent care of the fish.

## Competing interests

The authors declare no competing or financial interest

## Author contributions

P. Herbomel conceived and supervised the project, and obtained the funding. M. Tauzin isolated the *cerise* mutant, gave it its name, and conducted the initial phenotypic study. D L Demy, M. Carrère and R. Noche conducted most of the subsequent study, with the help of M. Le Bris and C. Baek during their respective Master internship; M. Yousfi contributed the 4C4/Pu.1 double immunostaining images. D L Demy performed the initial mapping of the mutation; I. Leshchiner & W. Goessling identified the slc7a7 mutation via deep sequencing of mutant vs. sibling genome pools. P. Herbomel and D L Demy wrote the paper.

## Funding

This work was supported by grants to P.H. from the European Commission through the FP6 “ZF-Models’ Integrated Project, from the Fondation pour la Recherche Médicale (“FRM 2012 team” #DEQ20120323714, “FRM 2016 team” #DEQ20160334881) and from the Laboratoire d’Excellence Revive (Investissement d’Avenir; ANR-10-LABX-73).

## Supplementary Figures

**Fig. S1.**
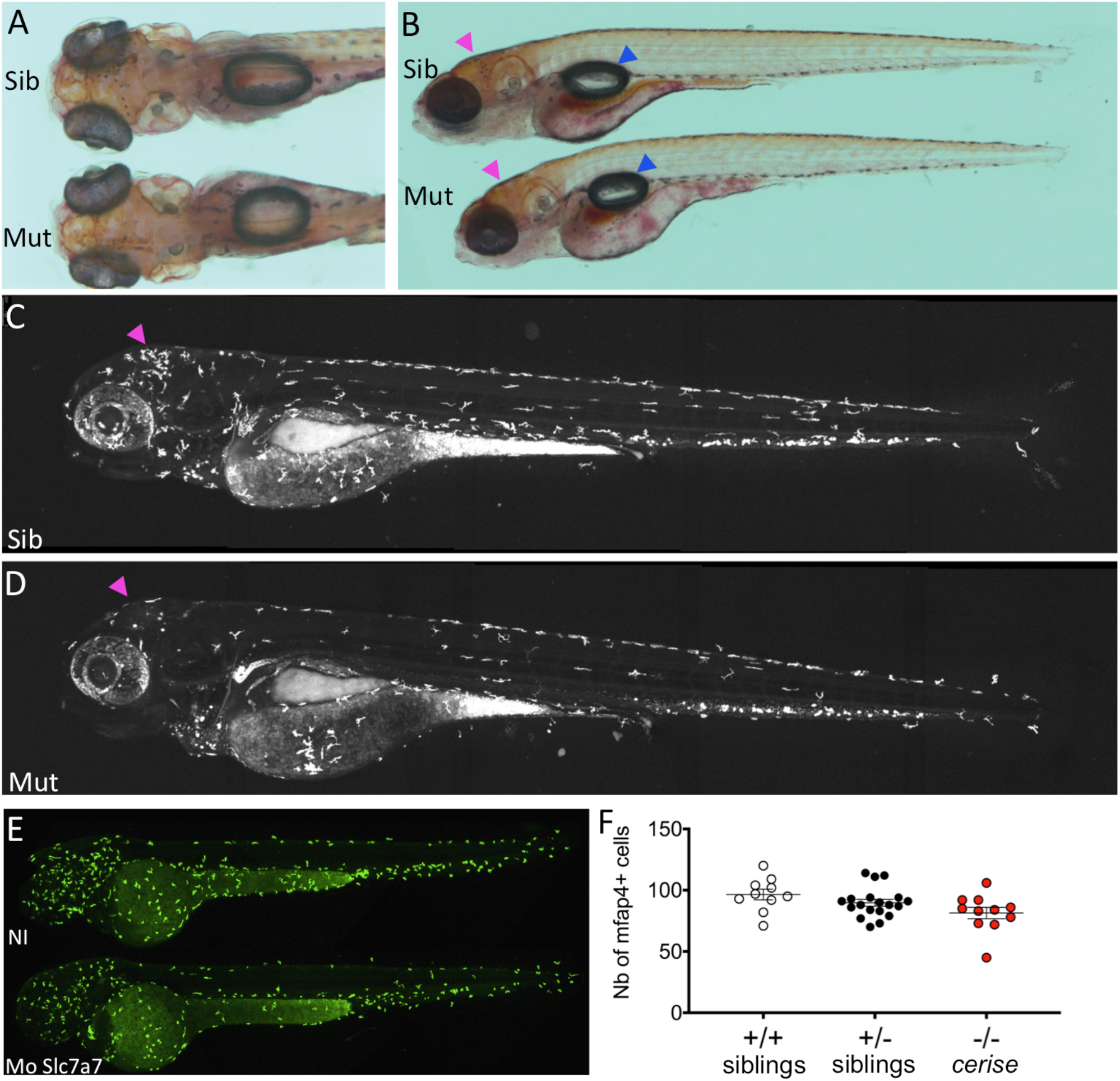
Primitive macrophages are normally produced in Slc7a7 deficient zebrafish. **(A-B)** Dorsal **(A)** and lateral **(B)** view of a live *cerise* mutant and a sibling stained with Neutral Red at 5 dpf. The mutant, recognizable by its absence of microglia in the optic tectum (pink arrowheads), has overall normal morphology, with a well developed swimbladder (blue arrowheads). **(C-D)** Lateral view of global macrophage population in live *Tg(mfap4:mCherryF)* control sibling **(C)** and cerise mutant **(D)** at 4 dpf. **(E)** Lateral view of global macrophage population in live *Tg(mpeg:Gal4;UAS:Kaede)* control or slc7a7-sp22 morphant embryos at 48 hpf. **(F)** Macrophage count in the whole body of 48 hpf *cerise* progeny according to their genotype, revealed by mfap4 ISH followed by qPCR genotyping. Pink arrowheads point to tectal microglia. Sib, sibling; mut, mutant; NI, non-injected; Mo, morpholino; Nb, number

**Fig. S2:**
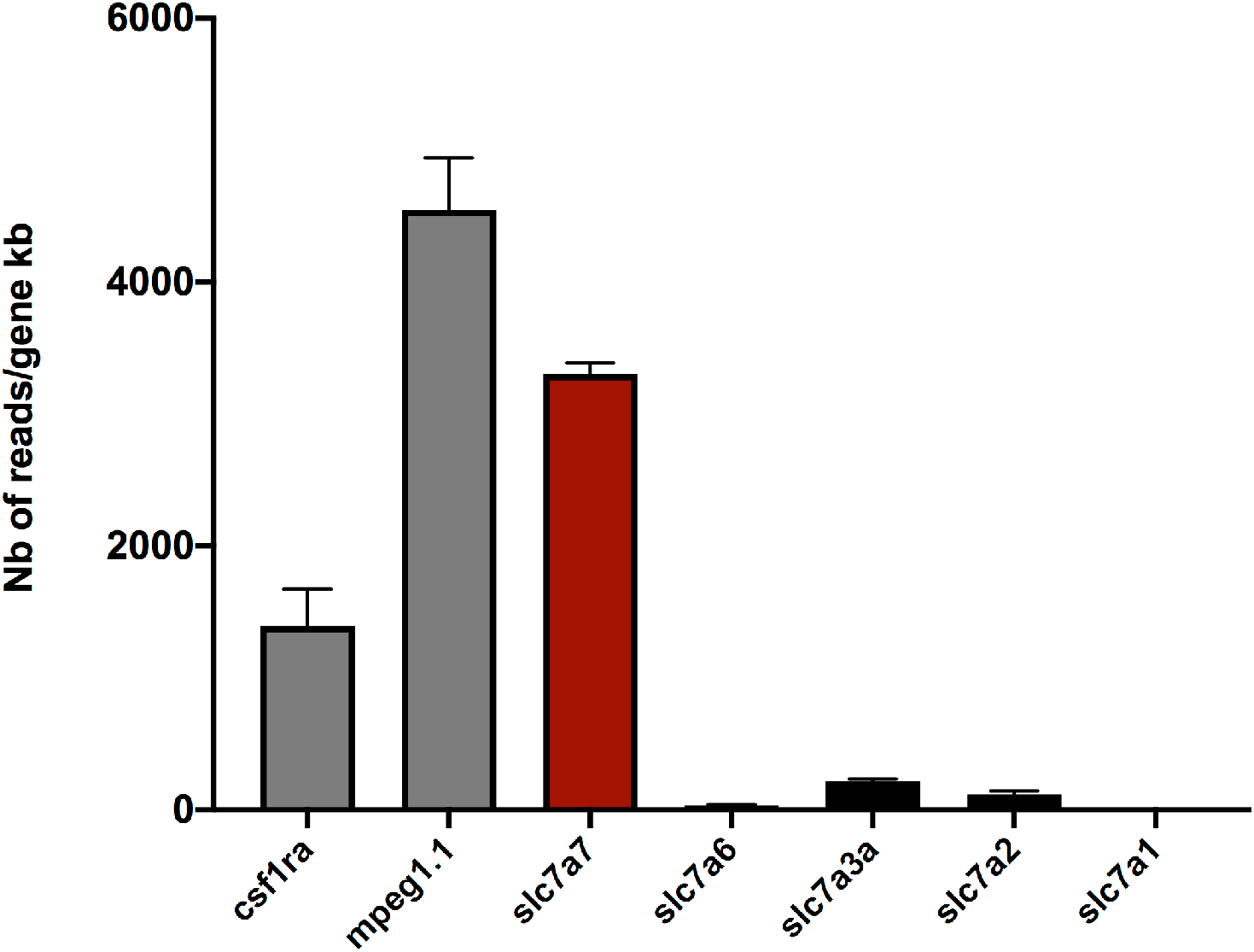
Slc7a7 is the most highly expressed cationic amino acid transporter in macrophages of zebrafish larvae at 3 dpf. Macrophages were FACS-sorted from WT zebrafish larvae at 3 dpf, and subjected to RNAseq analysis (see M&M section). Expression levels are given here as number of reads per kilobase of mRNA sequence, for two typical macrophage-specific genes (*csfr1a* and *mpeg1.1*), and the 5 cationic amino acid transporters of the slc7 family. See also Table 1 for raw data and additional genes.

**Fig. S3.**
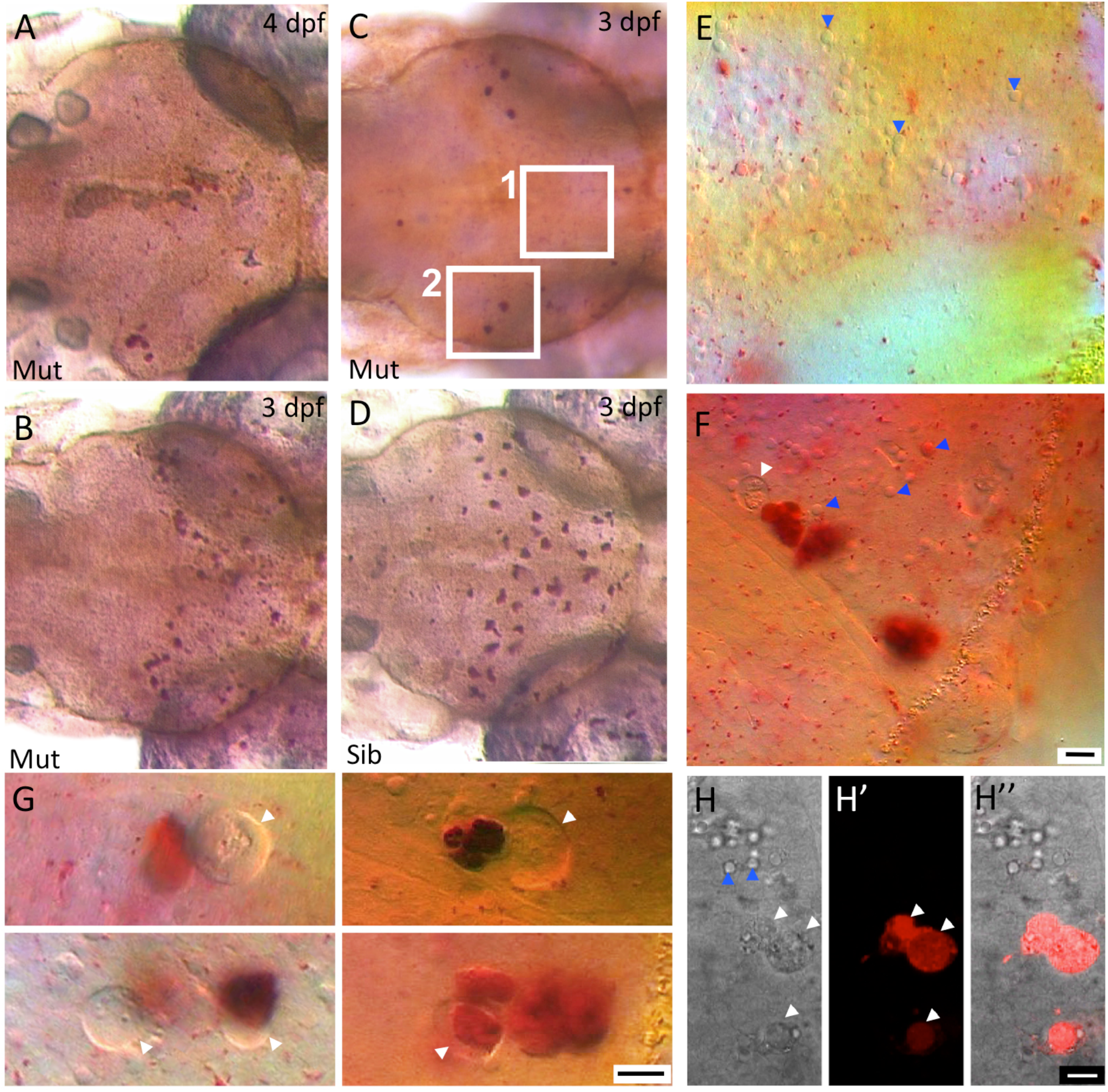
Close examination of neutral red stained *cerise* mutant larvae suggest that primitive macrophages are able to initiate brain colonization, but die soon thereafter. **(A-F)** Dorsal view of the optic tectum of live zebrafish larvae stained with Neutral Red; *cerise* mutants at 4 dpf **(A)** and 3 dpf **(B, C)**, sibling at 3 dpf **(D)**, and close-ups on the mutant NR+ material at 3 dpf **(E, F**) on regions shown by white frames in (C), and at still higher magnification in G. **(H-H’’)** Close-up on tectal microglia in live *Tg[mfap4:mCherryF] cerise* mutant larvae at 3 dpf. Blue arrowheads point to neuronal apoptotic bodies; white arrowheads point to abnormal large vacuoles (note that the DIC shadow cast contrast delineating their contour is inverted relative to that shown by the apoptotic bodies, indicating a lower refraction index, ie a less dense / more aqueous content) only in the mutant, always next to or including NR+ material. Scale bar, 10 μm. Sib, sibling; mut, mutant.

**Fig. S4.**
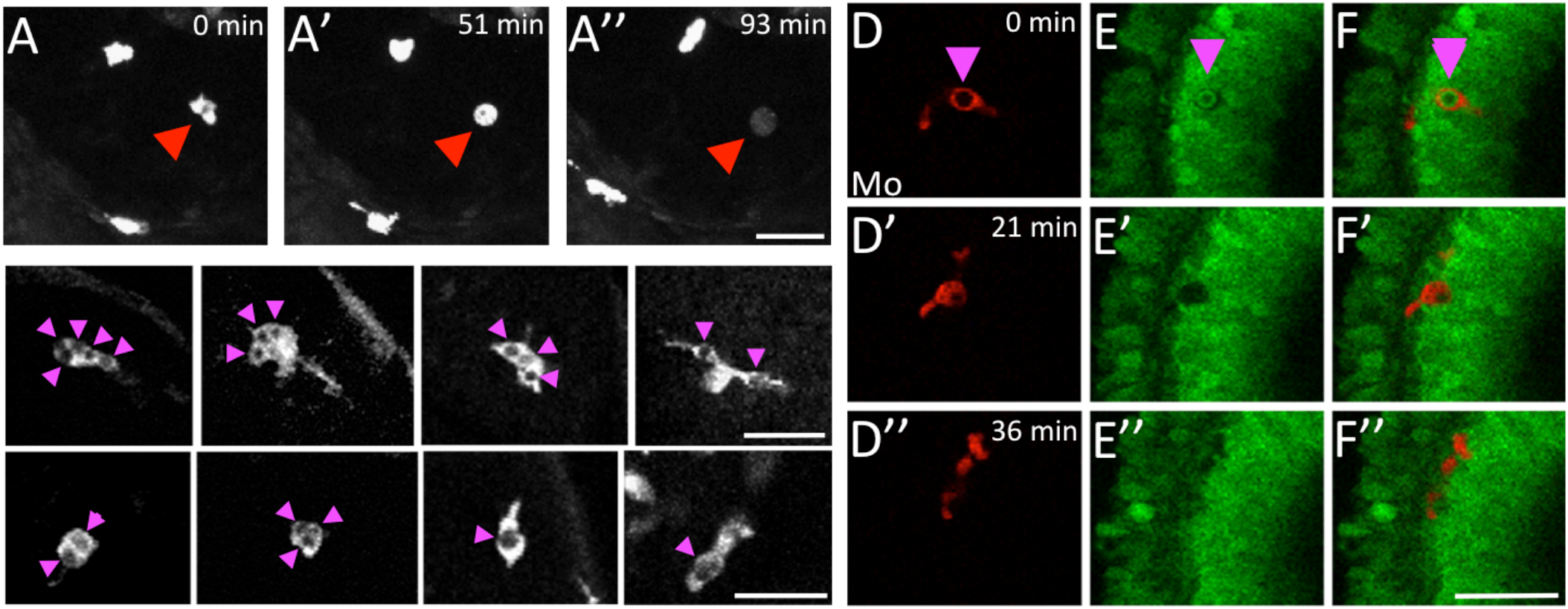
The death of microglia in *cerise* correlates with their efferocytic activity. **(A-A’’)** Close-ups on a dying mCherry+ macrophage in the retina of a 2.5 dpf *Tg(mpeg:Gal4; UAS:NfsB:mCherry)* slc7a7-sp22 morphant over time. See also Movie 2. **(B, C)** Close-ups on mCherry+ macrophages in the optic tectum of live *Tg(mpeg1:Gal4; UAS:NfsB-mCherry)* control (B) or slc7a7-sp22 morphant (C) larvae at 3 dpf. **(D-F’’)** Close-ups on an apoptotic HuC:GFP+ neuron (E-E’’) being engulfed by a mpeg1:mCherry+ macrophage which then dies (D-D’’; F-F’’), in the retina of a sp22 morphant over time. Scale bars, 30 μm. Pink arrowheads point to phagosomes inside macrophages and red arrowheads to dying macrophages. NI, non-injected; Mo, morphant

**Fig. S5.**
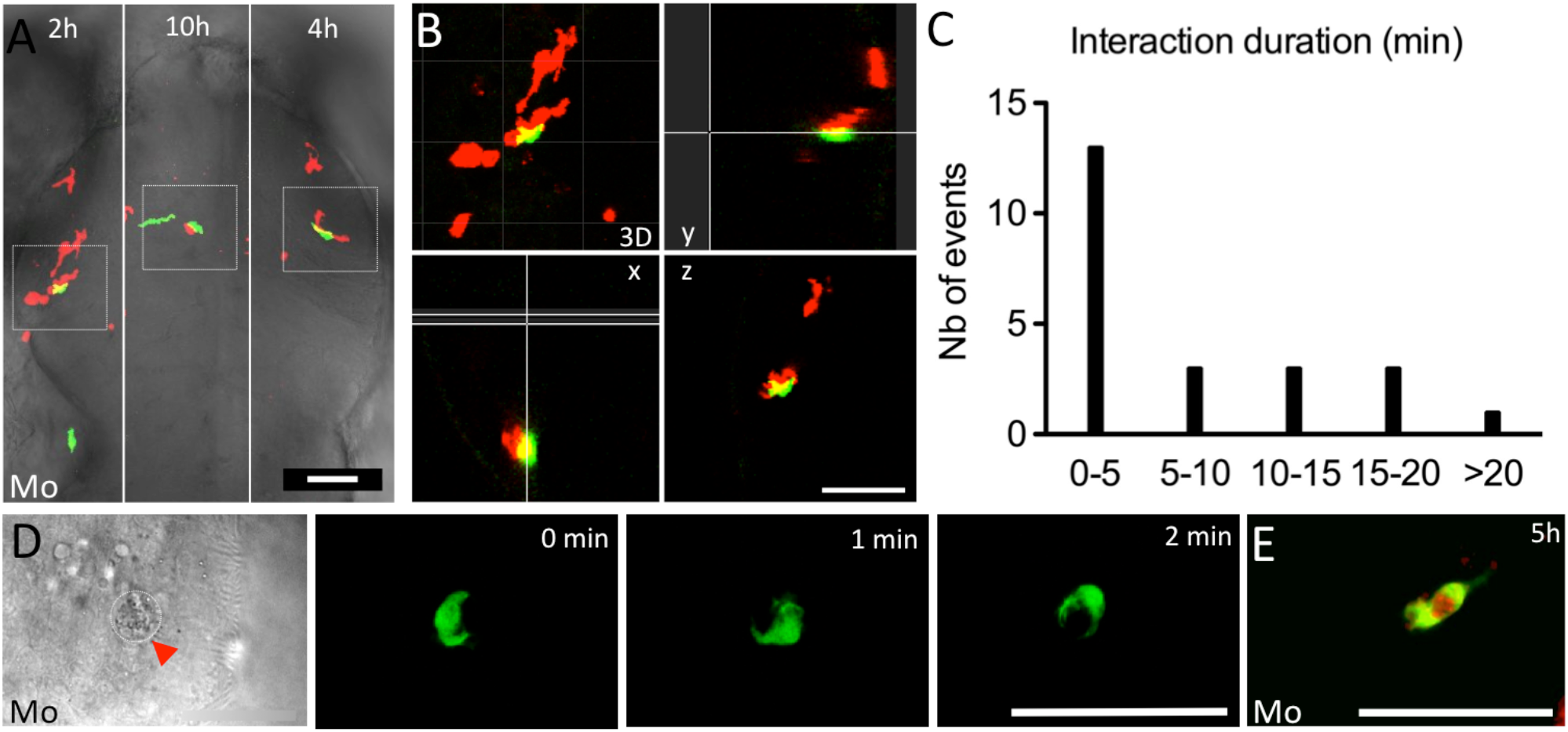
Neutrophil - macrophage/microglia interactions in the brain of *cerise* mutants. **(A, B, D, E)** Selected time points and image sub-fields of *in vivo* time-lapse confocal imaging sequences of mCherry+ macrophages/microglia and GFP+ neutrophils in the optic tectum of a slc7a7-sp22 morphant (A, B) or *cerise* mutant (D, E) at 3 dpf in dorsal view. **(A)** shows the whole tectum divided in three subfields (left, central, right) to display microglia / neutrophil interactions that occurred at different time points in each sub-field. **(B)** Close-up and orthogonal views of the interaction shown in the left sub-field (t=2h) in (A). **(C)** Distribution of the macrophage/neutrophil interaction durations. **(D, E)** Longer interactions observed in morphant larvae displaying only a few macrophage/microglia remnants in the tectum; (D) a neutrophil rapidly moving around a macrophage remnant and sticking to it for hours; (E) a neutrophil that engulfed mCherry+ material, presumably from a macrophage. Scale bars, 50 μm

**Fig. S6.**
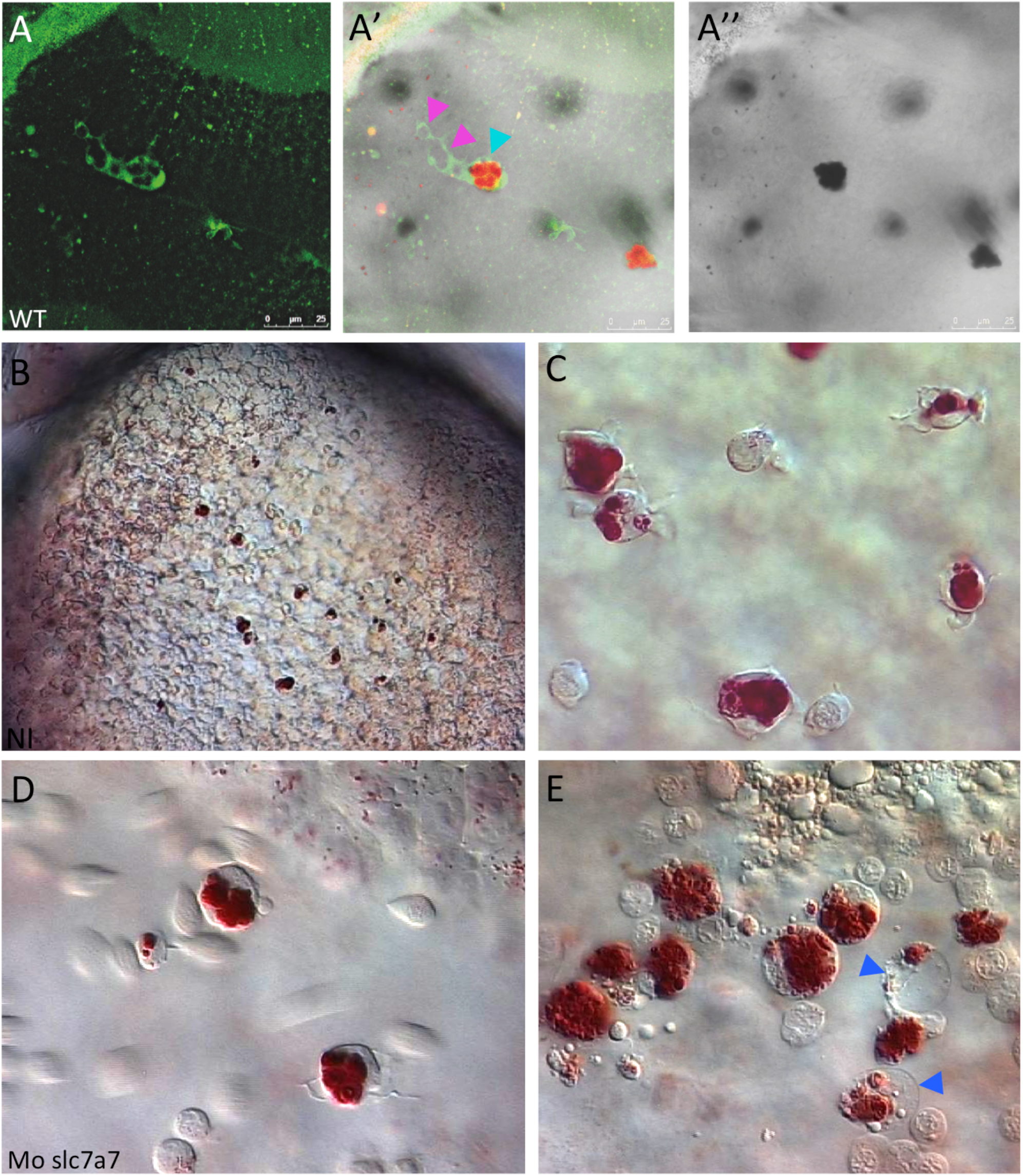
Vital staining of zebrafish macrophages with Neutral Red highlights their acidified effero-phagosomes. **(A-A’’)** Close-up on two mpeg1:GFP+ tectal microglial cells after Neutral Red vital staining at 4 dpf; (A), confocal fluorescence image; (A’), fluorescence + brigh-field transmitted light image; (A’’), bright-field only. Neutral Red accumulates in the already acidified effero-phagosomes (blue arrowhead), whereas more recent efferophagosomes are NR-negative (pink arrowheads). **(B-E)** VE-DIC/Nomarski observation in the yolk sac circulation valley (duct of Cuvier) of neutral red stained WT **(B, C)** and slc7a7 morphant **(D, E)** embryos at 30 hpf. While macrophages in slc7a7 morphants are as efferocytic towards apoptotic erythroblasts as in control embryos (C, D), some of them subsequently die, becoming a large empty vesicle containing particles in Brownian motion, beside the NR+ material that they previously engulfed (E, blue arrowheads); see also Movies 6 and 7.

## MOVIE LEGENDS

**Movie 1 (related to Fig. 2A-C). Dynamics of primitive microglia in the midbrain optic tectum of WT zebrafish larva by 3 dpf.** Time-lapse confocal imaging of a 3 dpf *Tg(mpeg1:gal4; uas:Nfsb-mCherry)* larva injected with a control morpholino at the 1-2 cell stage. Dorsal view (maximum projection from a z-stack); time step = 3 min.

**Movie 2 (related to Fig. 2A-C). Primitive microglial cells die in the midbrain optic tectum of a Slc7a7-deficient zebrafish larva by 3 dpf.** Time-lapse confocal imaging of a 3 dpf *Tg(mpeg1:gal4; uas:Nfsb-mCherry)* larva injected with the slc7a7-sp22 morpholino at the 1-2 cell stage, and imaged in parallel with the control Mo-injected larva shown in Movie 1, in the same glass-bottomed dish. Dorsal view (maximum projection from a z-stack); time step = 3 min. Selected time points of this movie are shown in Fig. 2C. The macrophages / microglial cells tracked by a white and a yellow arrowhead correspond to the dark and light green arrowheads in Fig. 2C, respectively. Another series of consecutive death/ engulfment/ death events occuring nearby in this movie is tracked by a blue/grey series of arrowheads in Fig. 2C. The z-stack used to produce the maximum projection shown here is somewhat thicker than in Fig. 2C, such that at the beginning, on the right side, it includes macrophages that are not within the brain parenchyme but above it.

**Movie 3 (related to fig. 2G-H’). Dynamics of primitive microglia in the retina of WT zebrafish larva from 2.5 dpf.** Time-lapse confocal imaging starting by 54 hpf of a *Tg(mpeg1:gal4; uas:Nfsb-mCherry)* larva injected with a control morpholino at the 1-2 cell stage. Lateral view (maximum projection from a z-stack); time step = 3 min.

**Movie 4 (related to fig. 2G-H’). Dynamics of primitive microglia in the retina of a Slc7a7-deficient zebrafish larva from 2.5 dpf.** Time-lapse confocal imaging starting by 54 hpf of a *Tg(mpeg1:gal4; uas:Nfsb-mCherry)* larva injected with the slc7a7-sp22 morpholino at the 1-2 cell stage, and imaged in parallel with the control Mo-injected larva shown in Movie 3, in the same glass-bottomed dish. Lateral view (maximum projection from a z-stack); time step = 3 min.

**Movie 5 (related to Fig. 3F). Neutrophils infiltrate the retina in Slc7a7-deficient larvae by 3 dpf.** Time-lapse confocal imaging of a *Tg(mpx:GFP)* 3 dpf larva injected with the slc7a7-sp22 Mo at the 1-2 cell stage. Lateral view of the eye (maximum projection from a z-stack); time step = 1 min.

**Movie 6 (related to Fig. S6). Neutral red vital staining of macrophages that engulfed numerous erythroblasts in the yolk sac of a WT embryo by 30 hpf.** Real-time VE-DIC microscopy. In all zebrafish embryos, a variable number of erythroblasts undergo apoptosis by 24-30 hpf and are engulfed by the primitive macrophages (Herbomel et al. 1999); the embryo shown here belonged to a clutch in which this rate of early erythroblast apoptosis was in the higher range.

**Movie 7 (related to Fig. S6). Neutral red vital staining of macrophages that engulfed numerous erythroblasts in the yolk sac of a Slc7a7-deficient embryo by 30 hpf.** Real-time VE-DIC microscopy. This embryo is from the same clutch and at the same stage as the control embryo shown in Movie 6, but was injected with the Slc7a7-sp22 Mo at the 1-2 cell stage. Its macrophages show a similar number of engulfed, neutral red stained erythroblast corpses, but unlike in the control embryo, some of these macrophages (two in the shown field, pointed at by blue arrowheads in Fig. S6E) died thereafter, as evidenced by their empty content but for particles in Brownian motion.

**Table S1.**
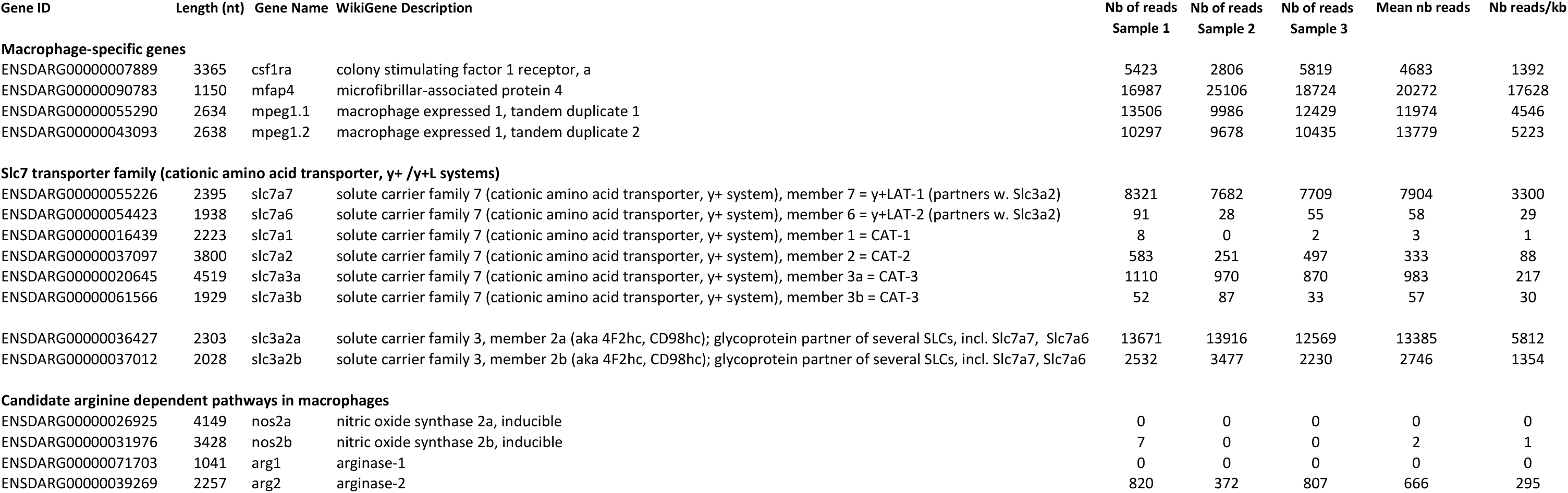
Gene expression levels of Slc7 family members and arginine-dependent enzymes in macrophages of zebrafish larvae at 3 dpf. Fluorescent macrophages were sorted from three different batches (replicates) of 3 dpf transgenic larvae (see M. & M.) and submitted to RNAseq analysis. 10 Mreads of 50 nt (0.5 Gb) were obtained for each replicate. The gene expression level is expressed in Number of reads per kilobase of RNA transcript.

## Notes

### Competing Interest Statement

The authors have declared no competing interest.

